# Functional Integration of Different-Sex Gonad Transplants into the Adult Mouse Hypothalamic Pituitary Gonadal Axis

**DOI:** 10.1101/2025.07.21.666020

**Authors:** Daniel R. Pfau, Monica A. Rionda, Evelyn Cho, Jamison G. Clark, Robin E. Kruger, Ruth K. Chan-Sui, Vasantha Padmanabhan, Molly B. Moravek, Ariella Shikanov

**Affiliations:** Obstetrics & Gynecology, University of Michigan, Ann Arbor, MI 48109; Pediatric Endocrinology, University of Michigan, Ann Arbor, MI 48109, US; Biomedical Engineering, University of Michigan, Ann Arbor, MI 48109, US; School of Social Work, University of Michigan, Ann Arbor, MI 48109, US; Molecular and Integrative Physiology, University of Michigan, Ann Arbor, MI 48109, US; Obstetrics, Gynecology and Reproductive Biology, Michigan State University, East Lansing, MI 48824, US; Reproductive Endocrinology and Infertility, Department of Women’s Heath, Henry Ford Health, Detroit, MI 48322, US; Cellular and Molecular Biology Department, University of Michigan, Ann Arbor, MI 48109, US

**Keywords:** Transgender, gender-affirming hormone therapy, testis, ovary, sex difference

## Abstract

Gender-affirming hormone therapy (GAHT) relies on exogenous hormones to produce hormonal milieus that achieve and/or maintain embodiment goals. Another potential route to these endpoints is transplantation of novel steroidogenic tissue. To develop a pre-clinical model, we asked whether different-sex gonad transplants can be functionally integrated into the adult mouse hypothalamic-pituitary-gonadal (HPG) axis. Adult male and female mice were gonadectomized and implanted with gonads from genetically matched but different-sex pups. Controls received gonads from same-sex pups. Temporal changes to gonadotropin and steroid hormone levels reveal the decoupling of the HPG following gonadectomy and gonad-dependent levels after transplanting donor gonads. After six weeks, histological structures in transplanted gonads were consistent with expected steroidogenesis and gametogenesis. Interestingly, pituitary, ARC and AVPV mRNA showed gonad- and sex-dependent expression patterns. Future work with this technique could lead to translation to gender affirming care and explorations of gonad-dependent sex differences in biomedical and basic research.

## Introduction

Many transgender, non-binary and gender diverse (TNG) individuals utilize GAHT to produce hormonal milieus that achieve and maintain embodiment goals^1^. Individuals may take testosterone (T) to increase circulating levels and/or suppress ovarian function, or estrogen (E), progesterone and/or anti-androgens (collectively termed E-GAHT) to increase their levels and suppress testicular function and androgen action^2,3^. Attaining physical results that cannot be generated by GAHT may involve gender-affirming surgeries^4^. While gonadectomies may remove the need to suppress gonadal function, continued GAHT use is required to maintain optimal steroid hormone levels for overall health^2^. The range of GAHT regimens and surgical procedures reflects variation in individuals’ responses to treatments and diverse patient needs^5–7^. As these needs are better understood^7,8^, novel interventions may increase the number of desirable phenotypes gender-affirming treatments can offer^9,10^. One potential surgical intervention could involve transplantation of different-sex donor gonads, which would rely on plasticity of hypothalamic and pituitary function to stimulate endogenous hormone production from transplanted gonads. Unlike GAHT, this procedure would reintroduce gonad cycles and homeostatic control of steroidogenesis, which may improve patient outcomes. For example, gonadectomy followed by T-GAHT is detrimental to bone mineral density measures for female mice but providing low-dose E, to mimic the low levels of circulating E produced by the testis, can rescue musculoskeletal architecture^11^. Hypothetically, the results achieved with pharmacological T-GAHT and E-GAHT could be reproduced and/or improved by endogenous testicular and ovarian steroidogenesis, respectively.

It has been well-established in humans and animal models that transplanted autologous ovarian tissue, removed for fertility preservation prior to gonadotoxic cancer therapies, restored ovarian endocrine function, demonstrated by elevated levels of ovarian hormones, regular menses and even live births^12–15^. Though relatively less studied, autologous transplants of testicular tissue or cells also restore hormone function in non-human primates^16,17^ and survived up to 6 months in the only human case study^18^. For these transplants, same-sex gonadal tissue integrates into the hosts hypothalamic-pituitary axis by responding to circulating gonadotropins and secreting hormones to reinstate the negative and positive feedback necessary for steroid hormone homeostasis and gonad function. Integration of transplanted different-sex gonads into the HPG axis of the host requires these three tissues communicate in a novel way. Though hypothalamic and pituitary sex differences enable the unique functions of each gonad^19–22^, the hypothalamus and pituitary of humans respond to circulating T and E levels regardless of an individual’s sex^23–26^. Further, ovarian tissue transplanted from female into male rhesus monkeys led to pre-ovulatory gonadotropin surges and cyclic gonadotropin and steroid hormone release^27^. Unlike ovaries, testicular transplants into different-sex non-human primates have not been performed. It is currently unknown whether human hypothalamic and pituitary plasticity extends to maintaining steroid hormone homeostasis and cycles for different-sex gonads.

Rodent models are being developed to inform and improve many gender-affirming treatments^28–35^. Like in human GAHT patients, the HPG axis of GAHT-treated mice responds to circulating T and E levels regardless of the animals sex^30,31^ suggesting they may be used to explore different-sex gonad transplant integration into a novel HPG^22^. In rodents, hypothalamic and pituitary sex differences are considered organized prior to adulthood^36,37^; however, new neurocircuitry to maintain gonad-specific HPG functions is integrated throughout adulthood^38^.

Since sex-dependent hormonal cues are known to mediate the integration of adult-born cells necessary for sex differences in HPG function^38^, it is possible that the circulating hormonal milieu produced by transplanted gonads can do the same. Here, we asked whether different-sex gonad transplants can be functionally integrated into the adult mouse HPG axis after a puberty driven by natal gonad type (see Figure 1 for study plan). Our objective was to determine the feasibility of different-sex gonad transplants in adult mice and whether mice could offer a useful model for further investigations of different-sex gonad transplants and gonad-dependent sex differences.

**Figure 1:**
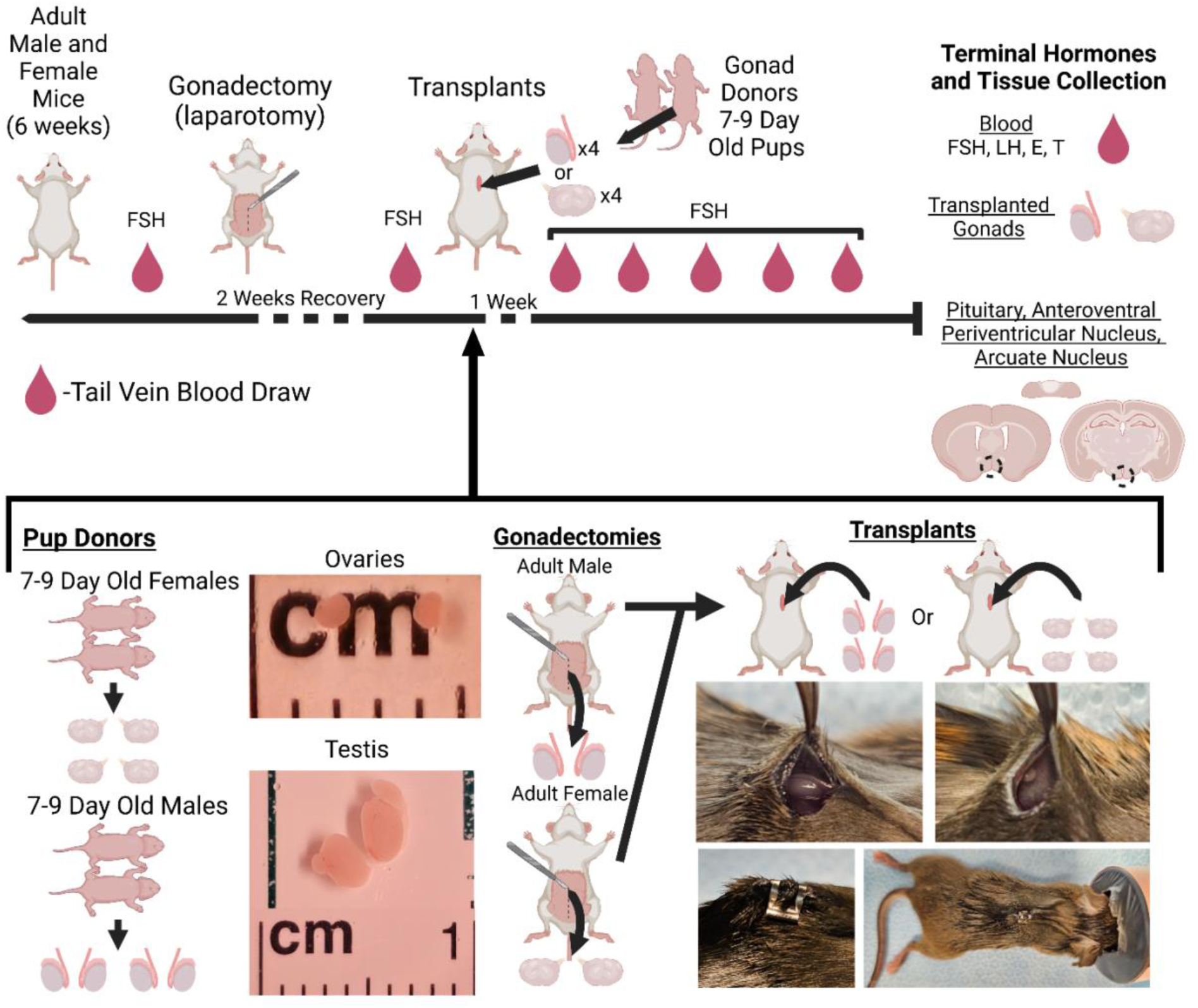
Study Design. E=Estradiol, T=Testosterone, FSH=Follicle stimulating hormone, LH=Luteinizing hormone. Created using Biorender.com.

## Methods

### Animals

Adult male and female mice (n=8/group) of the B6CBAF1/J (Jackson Laboratories) strain were co-housed five in a cage (L:D, 12:12) and provided food and water ad libitum throughout the experiment. Donor B6CBAF1/J pups (PD 6-9) were generated from our colony of C57BL/6J and CBA/J breeding pairs (Jackson Laboratories), held under the same conditions. All animal procedures were approved by the Institutional Animal Care & Use Committee at the University of Michigan (PRO00009635).

### Blood Collection and Surgical Procedures

Adult animals were left undisturbed for one week then blood serum was collected from tail veins the next week to measure follicle stimulating hormone (FSH) levels. The week after blood collection, all animals were gonadectomized via laparotomies then given two weeks to recover before collecting another blood sample to measure FSH. One week later, cages were randomly assigned to receive transplanted gonads from two different- or same-sex donor pups, for a total of four transplanted gonads (Fig 1). Young donors were used to maximize the number of implanted primordial/primary follicles and stem Leydig cells per unit volume of the tissue. Ovaries from 6-9 days old mice contain mostly primordial and a small fraction of activated primary and secondary follicles, while ovaries from adult mice contain corpus lutea and antral follicles^39^. Furthermore, human ovarian cortex would be used in a clinical setting and, like prepubertal mouse ovaries, contain mostly primordial follicles. Similarly, testis from 6-9 days old mice contain mostly stem Leydig cells^40^ and, pre-pubertal tissue or stem Leydig cells are used for human and non-human primate same-sex testis transplants^16–18^. After a week of recovery, 5 weekly tail vein blood draws were taken to measure FSH, followed by terminal cardiac blood collected on week 6 to analyze E, T, FSH and luteinizing hormone (LH).

### Terminal Measures

Animals were weighed and tissues sensitive to T or E levels measured, including clitoral area and uterine and seminal vesicle weight. Transplants were removed, imaged, and a portion placed in Bouin’s fixative. The brain was removed, and the pituitary placed in RNA protect. Brain tissue collection took place in sterile saline chilled over blue ice using sterile materials cleaned with RNAse Away (Thermo Scientific) per manufacturer instructions. The brain was placed in a brain matrix (Electron Microscopy) then a razor blade inserted directly posterior to the optic chiasm. Two more razor blade were placed 1 mm and 2 mm anterior to the first then a fourth 2 mm posterior to the first. The anterior and posterior brain slices were placed on a slide then a blunt hypodermic needle used to biopsy the AVPV and ARC before placing them in RNA protect.

### Hormone Analysis

Serum samples were kept at −20C before being shipped to the Ligand Assay and Analysis Core Facility, University of Virginia Center for Research in Reproduction. Sensitivity for each assay were as follows: Follicle Stimulating Hormone (sensitivity: 3 ng/mL, inter-assay CV 9.4%; In house radioimmunoassay^41^), Testosterone (sensitivity: 10 ng/dL, inter-assay CV 9.6%; IBL ELISA, Minneapolis, MN), Estradiol (sensitivity: 5 pg/mL, inter-assay CV 10.2%; ALPCO ELISA, Salem, NH), Luteinizing Hormone (sensitivity: 0.04 ng/mL, inter-assay CV 6%; In house radioimmunoassay^42^).

### Histology

After 24 hours of Bouin’s fixation, gonad transplant portions were washed in ethanol dilutions, stored in 70% ethanol, then embedded in paraffin blocks by the University of Michigan Dental School Histology Core. Transplant tissue, potentially containing multiple transplanted gonads, were sectioned serially at 5 um, 5 sections to a slide. Every other slide was stained with hematoxylin (Epredia, Kalamazoo, MI) and eosin (Ricca Chemical, Arlington, TX; H&E), then imaged at 5X. Morphological features were evaluated by tracking structures through Z-stacks using ImageJ, including seminiferous tubules, follicles, and corpora lutea (CL).

### Immunohistochemistry

Leydig cells and CLs were identified in H&E-stained testicular and ovarian tissue, respectively, then adjacent sections were used to visualize LH receptor location using a previously published paradigm^43^. One section from one slide was stained for each animal. All procedures were performed at RT and in tris-buffered saline (TBS) unless noted otherwise. Tissue mounted on slides were deparaffinized with xylenes, dehydrated, blocked with 0.3% hydrogen peroxide, and incubated in 0.1M citrate buffer (pH=6.0, Thermo Fisher, AAJ63950AP) for 20 minutes at 90°C then cooled to RT for 20 minutes. Tissue was permeabilized and blocked with 10% normal goat serum (NGS, Abcam, G9023) in TBS with triton for one hour. Sections were left overnight at 4°C with rabbit anti-LHCGR (Bioss, 1:250, bs-0984R) in 1% NGS then incubated with biotinylated goat anti-rabbit secondary (Abcam, AB64256) for one hour. Following streptavidin horseradish peroxidase (Abcam, AB64269) the sections were incubated in the DAB reaction (Abcam, AB64238) for 8 minutes before being counterstained with hematoxylin.

### RNA Extraction and qPCR

A RNeasy Kit (Qiagen) was used to extract RNA from brain and pituitary tissue following manufacturer procedures. RNA concentrations were measured using a NanoDrop (Thermo Fisher) before converting equal amounts to cDNA using an iScript cDNA Synthesis Kit (Bio-Rad, Hercules, CA). Target genes included *Esr1* and *Ar* for all tissues, *Kiss1, Gpr54, and Pgr* for the AVPV, *Kiss1, Gpr54, Pdyn, Tac2, and Npy* for the ARC, and *Cga, Fshb, Lhb, and Gnrhr* for the pituitary. *Fshb* Housekeeping genes tested included *Gapdh, Ppia, Rpl37* and *Sdha* (see Table S1 for primers; Integrated DNA Technologies, Coralville, IA). *Ppia* was chosen as the housekeeping gene for the ARC and AVPV, and *Gapdh* for the pituitary. An iTaq Universal SYBR green Supermix (Bio-Rad) was used for reactions that were run using a roto-gene Q (Qiagen) system. LinRegPCR software^44^ was used to calculate fold change values.

### Statistical Analysis

Planned comparisons were used to analyze differences in pre- and post-gonadectomy FSH levels; pre-gonadectomy differences between sexes, within-sex levels before and after gonadectomy, and the differences in FSH levels pre/post-gonadectomy between sexes were analyzed using unpaired Mann-Whitney or paired Wilcoxon tests (alpha=0.017). Normality for all analyses was determined using the Shapiro-Wilkes test. Uterine and seminal vesicles weights and clitoral areas were compared with t-tests (alpha=0.05). A two-way ANOVA with animal sex and gonad type as factors was performed for all other measures followed by posthoc pairwise comparisons adjusted for multiple comparisons using Tukey’s method, to investigate significant main effects or interactions (alpha=0.05). Data for 2-way ANOVA analysis without normal distributions were log transformed in Graphpad/PRISM before analysis. Outliers were visually assessed and removed from analysis if 1.5 times the interquartile range above the third or below the first quartile (see Table S3,S4).

## Results

### Terminal Gonadotropin and Steroid Hormone Levels are Gonad-Dependent

FSH levels are higher pre-gonadectomy in male than female mice (Fig 2A; See Table S2 for hormone Means±*SD*; see Table S3,S4 for individual values; *p*<0.0001, *d*=3.8). Following gonadectomy, an increase in circulating FSH was evident in males and females (Males: p=0.0002, *d*=1.3; Females: p<0.0001, *d*=6.3), but the increase was greater in females (*p*<0.0001, *d*=2.7, Pre: 4.2±2.7 ng/mL FSH, Post: 38.6±3.8, Difference: 34.4±3.3) compared with males (Pre: 38.1±8 ng/mL FSH, Post: 50.6±5, Difference: 12.46±6.5). Terminal FSH and LH levels were higher in animals with testes (Fig 2B,C; FSH *p*<0.0001, *d*=1.3; LH *p*=0.0003, *d*=1.1) but independent of sex (FSH *p*=0.9, *d*=0.03; LH *p*=0.07, *d*=0.2). Terminal E levels were independent of both gonad type (Fig 2D, *p*=0.4, *d*=0.08) and sex (*p*=0.1, *d*=0.1). Mice with testes had higher terminal T levels than those with ovaries (Fig 2E, *p*<0.0001, *d*=1.2), independent of sex (*p*=0.7, *d*=0.2).

**Figure 2:**
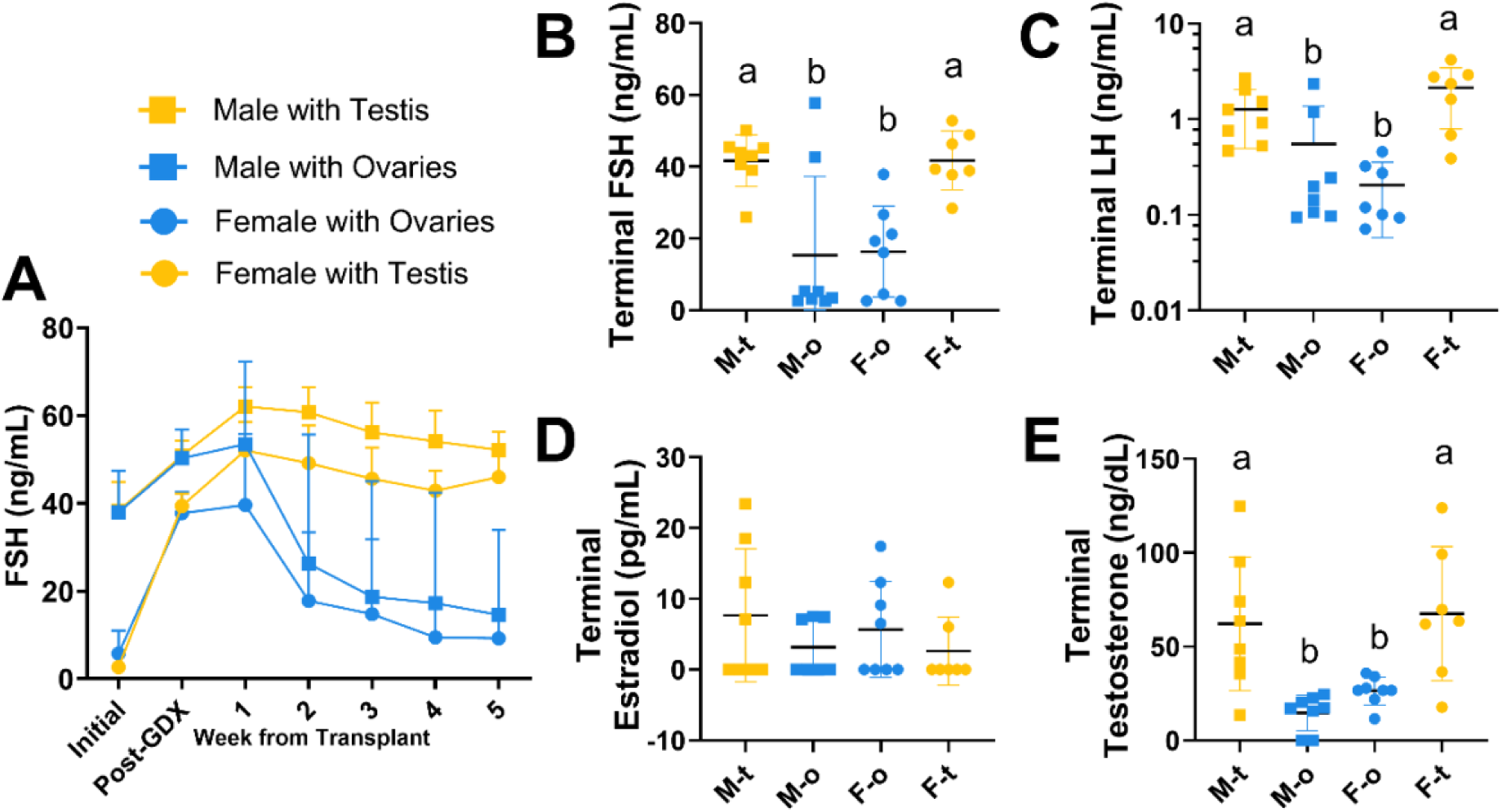
Weekly and terminal hormones levels. A) Follicle stimulating hormone (FSH) levels were higher in males compared with females pre-gonadectomy. FSH was elevated in both males and females post-gonadectomy then the average starts lowering two weeks after gonad transplants were performed. B) Terminal FSH levels were highest in animals with testes. C) Likewise, terminal luteinizing hormone was highest in animals with testes. D) Terminal estradiol levels were not gonad- or sex-dependent while E) terminal testosterone was higher in animals with testes compared to those with ovaries. Different letters are statistically different, *p*<0.05

### Anatomical Changes Are Gonad-Dependent

Male mice weighed more than females (see Table S3,4 for individual values, *p*<0.0001, *d*=1.5) regardless of gonad type (p=0.11, *d*=0.08). Female mice with testes had larger clitorises than those with ovaries (see Table S3,4 for individual values, *p*=0.004, *d*=1.7) while female mice with ovaries had heavier uteri than those with testes (Fig 3A,B, *p*=0.014, *d*=1.1). Male mice with testes had heavier seminal vesicles than males with ovaries (Fig 3C,D; *p*=0.002, *d*=2.7).

**Figure 3:**
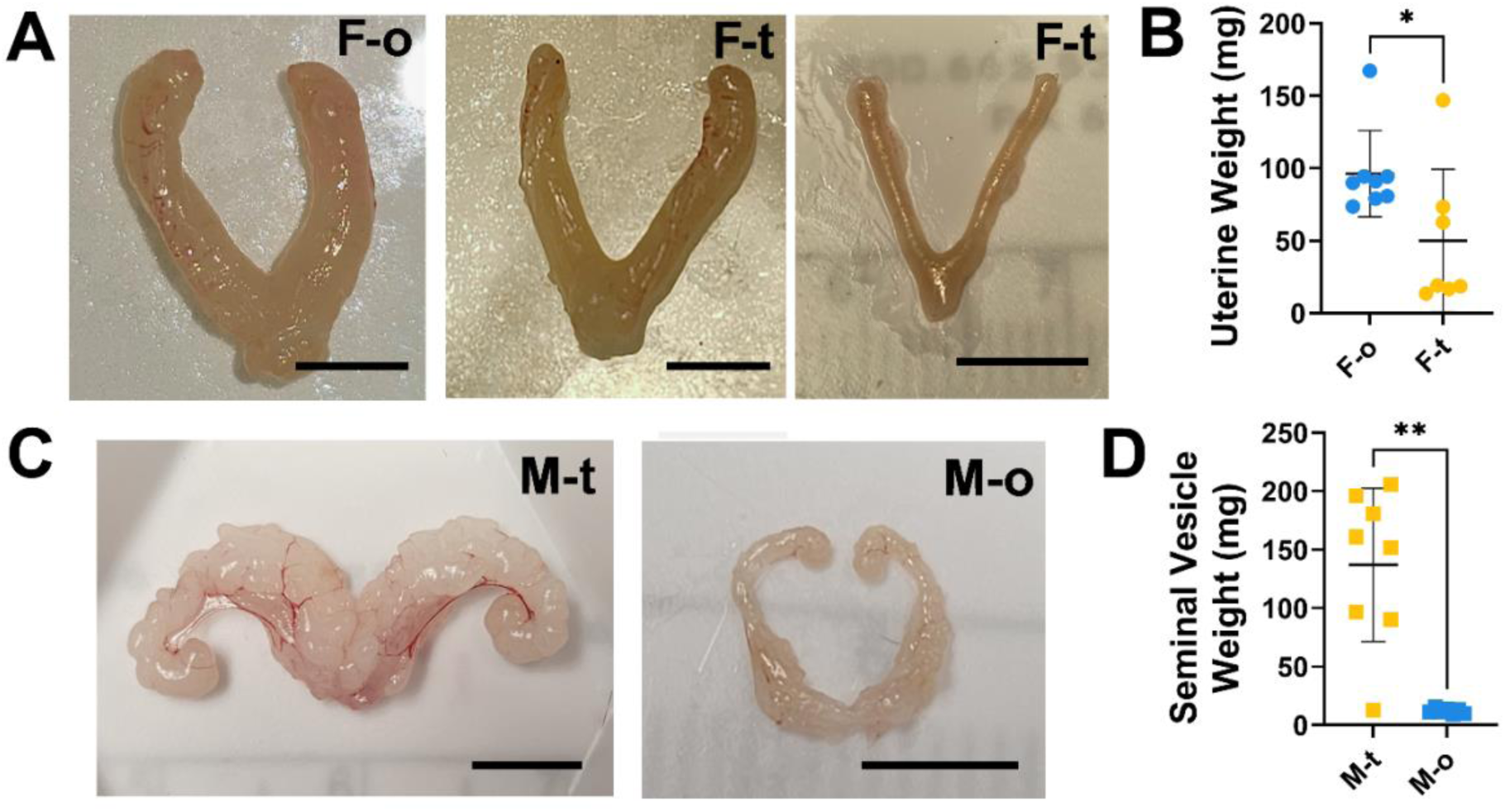
Reproductive Anatomy. A) Uterus B) weight was highest in females given ovary (F-o) transplants and there was large variability in the size of uteri from females implanted with testes (F-t). C) Seminal vesicles were D) heavier in males that received testes (M-t) transplants compared with those given ovaries (M-o). Different letters are statistically different, *p*<0.05.

### Testis Transplant Anatomy and Histology Indicate Successful HPG Integration

Transplanted testes fused to form connected structures, likely containing multiple transplanted gonads (see Table S3; Fig 4A,B,E,F,I,J,L), and embedded in muscular/adipose (Fig 4A,C) and epidermal tissue (Fig 4B,D,E,G,K). Before removal, vasculature was visible, surrounding testis transplants (Fig 4A,B,C,D,G,I,K). Most testis transplants displayed irregular margins but several transplants removed from female hosts were smooth (Fig 4C,D) or had what appeared to be unhealthy tissue (Fig 4H,K). Histology revealed structures resembling seminiferous tubules were present in all testis transplants (Fig 4). All testis transplants, aside from one taken from a female, had seminiferous tubules with varying degrees of spermatogenesis up to the spermatocyte stage (Fig 4M). These tubules displayed spermatogonia (Arrowheads), spermatocytes (Arrows), and cytoplasmic processes from Sertoli cells extending into their lumens (Fig 4M). Some structures resembling seminiferous tubules lacked these cytoplasmic processes and had empty lumens (Fig 4N). No testis transplants had recognizable spermatids. Leydig cells (Chevron) were identified in all testis transplants taken from male and female mice, based on their morphology, location between the seminiferous tubules (Fig 4O), and the presence of LH receptors (Fig 4P).

**Figure 4:**
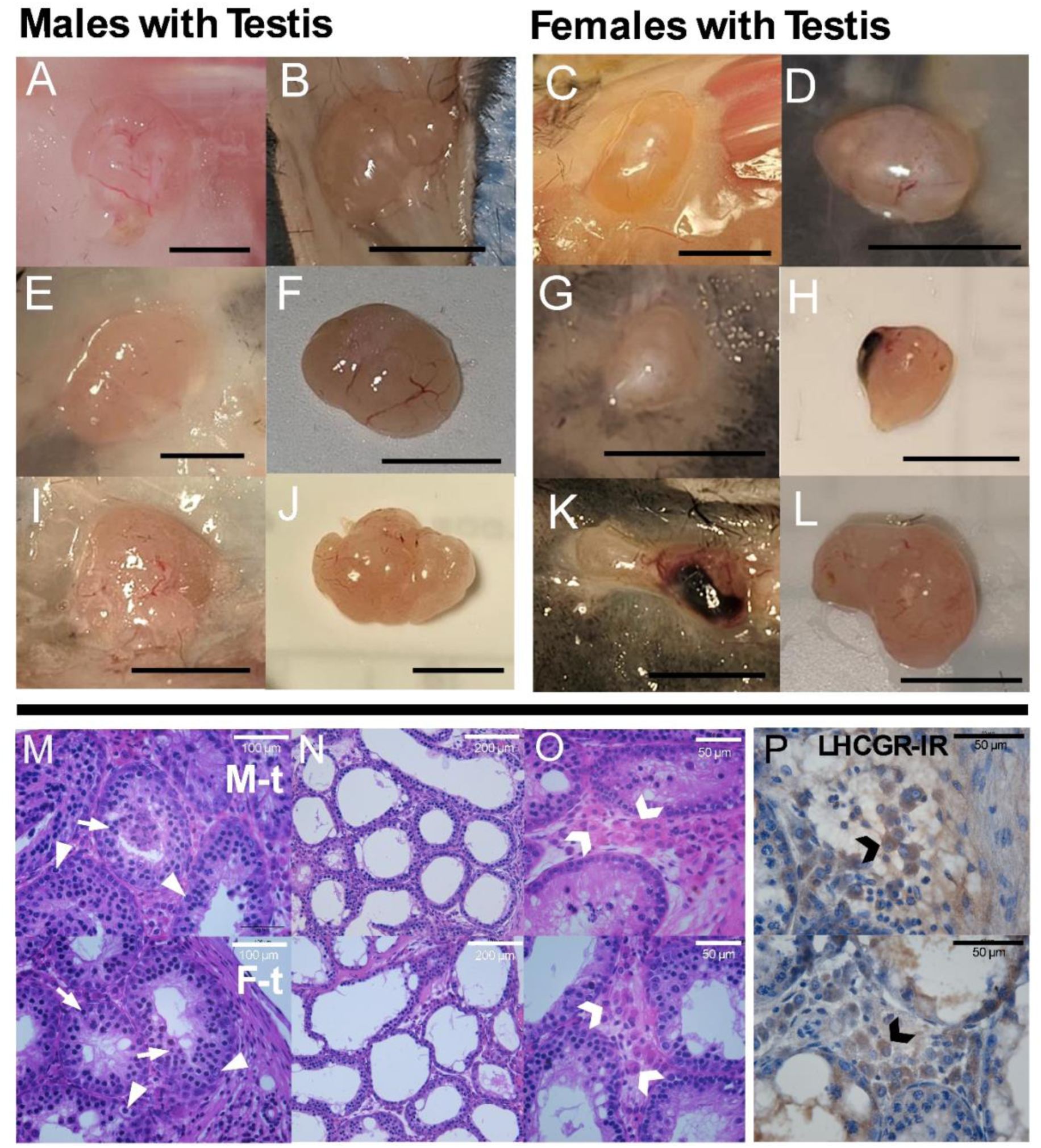
Representative images of transplanted testes anatomy and histology. A-L) testes transplants from male (M-t) and female (F-t) animals display variable growth, vasculature and morphology. Transplants were attached to the epidermal layer or embedded in muscle/adipose tissue. There is variability in overall transplant size and the presence of irregular/smooth margins or bloody portions, which may indicate differences in the number of donor gonads that survived and grew. M) Histology indicates seminiferous tubules (hematoxylin and eosin) are present in all transplants with clear evidence of spermatogonia (arrowhead) and primary spermatocytes (arrow). N) Many transplants contained seminiferous tubules with dilated lumens and thinner tubes. O) Leydig cells (chevron) were identified by their location between seminiferous tubules, shape, and P) expression of the luteinizing hormone receptor (Brown LHCGR-IR, blue hematoxylin counterstain).

### Ovary Transplant Anatomy and Histology Indicate Successful HPG Integration

Many ovarian transplants fused to form a structures containing multiple transplanted gonads in female and male mice (see Table S4; Fig 5A,B,D,F). Other ovarian transplants from female and male mice had multiple discrete portions that were, nonetheless, in close apposition (Fig 5C,I). Additionally, most ovary transplants had irregular margins (Fig 5A,B,D,F,H) with some protruding structures having a bloody appearance (Fig 5) and others resembling the opaque fluid-filled antrum of antral follicles (Fig 5B,D,F,H) or yellowed, potentially luteal, tissue (Fig 5A,C,F,K). Most transplanted ovaries from males and some from females had small bloody protrusions or patches (Fig 5E,G,I,J,K,L). Histological analysis of ovary transplants (Fig 5M) uncovered primary, secondary and tertiary follicles in all transplants, and most also had antral (Fig 5N) and atretic antral follicles, characterized by direct contact between oocyte (O) and antral fluid (A). However, one males’ ovary transplant contained only pre-antral follicles and cystic antral follicles. Three transplants taken from female mice had cystic antral follicles while these blood-filled structures were present in all ovaries taken from males (Fig 5M, asterisk). Corpora lutea (CL), identified by their histological structure (Fig 5O) and LHCGR receptor expression (Fig 5P), were present in all ovary transplants taken from female mice and in two taken from male mice.

**Figure 5:**
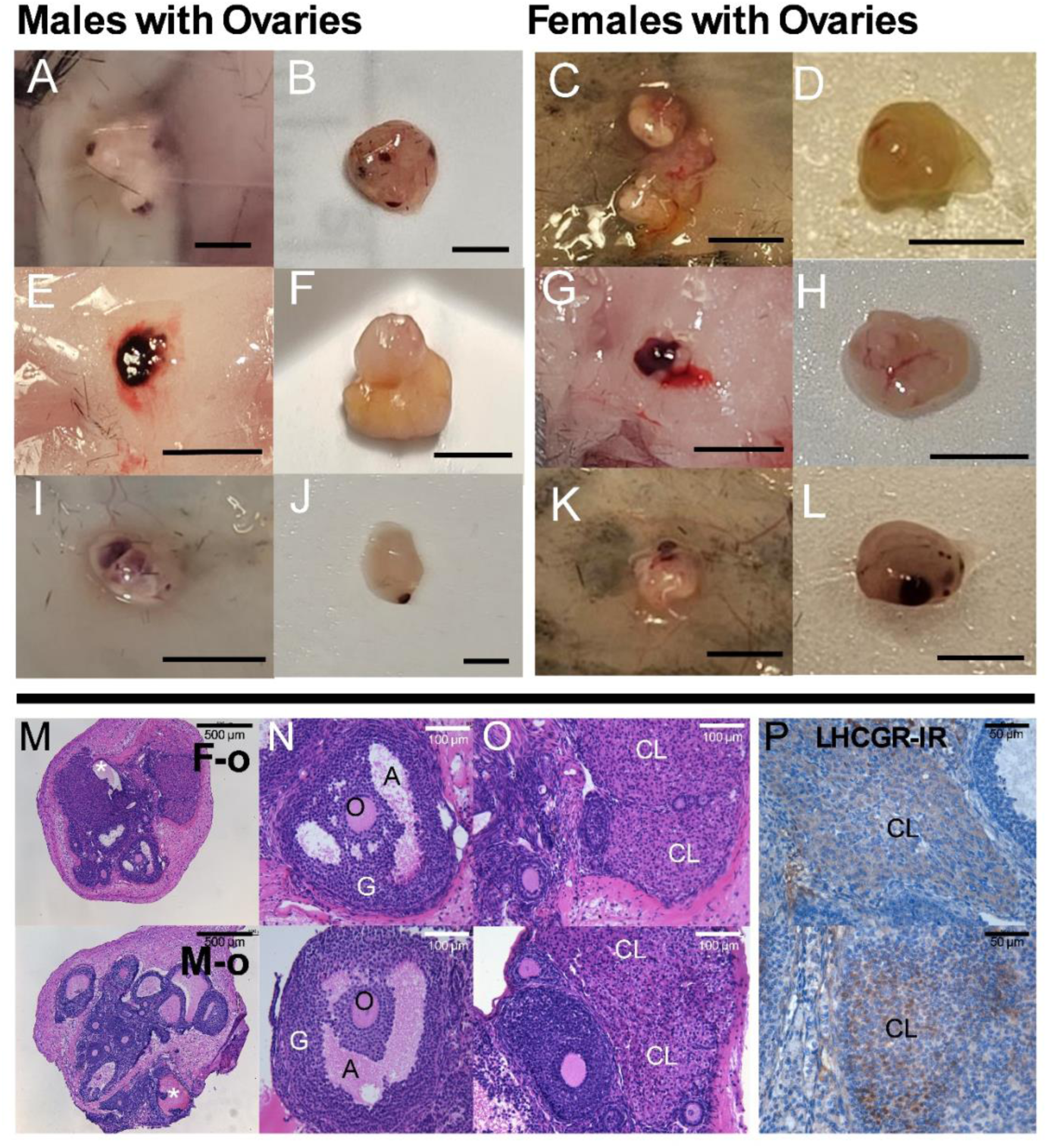
Representative images of transplanted ovary anatomy and histology. A-L) Ovary transplants from males (M-o) and females (F-o) show growth and vascularization. Many transplants taken from males, and a few from females, had small blood spots or larger blood patches, potentially sites of un-ovulated cystic follicles (asterisk). Structures resembling fluid-filled antral follicles and yellowed luteal tissue were present in ovaries from males and females. M) Ovarian histology (hematoxylin and eosin) indicates the presence of N) antral follicles (A) with intact antrum (A), oocytes (O) and granulosa (G) cells, and O) corpora lutea (CL), P) which express the luteinizing hormone receptor (Brown LHCGR-IR, blue hematoxylin).

### Hypothalamic and Pituitary Gene Expression is Sex- and Gonad-Dependent

Several genes across the HPG showed significant differences based on sex and/or gonad type (Fig 6). Within the AVPV, *Esr1* expression was highest in animals with ovaries compared to those with testes (Fig 6A; *p*=0.03, *d*=0.7) while AVPV *Kiss1* levels were higher in females than males (Fig 6B; *p*<0.001, *d*=1). There were no significant main effects of sex or gonad nor their interaction for the expression of AVPV *Ar*, *Gpr54*, and *Pgr* (Fig 6L). In the ARC, *Esr1* was highest in female animals compared to males (Fig 6C; *p*=0.01, *d*=0.7). Conversely, ARC *Tac2* (Fig 6D; *p*=0.03, *d*=0.6) and *Npy* (*p*=0.04, *d*=0.6) expression levels were higher in animals with testes than those with ovaries, regardless of sex. ARC genes without significant differences included *Ar*, *Kiss1*, *Gpr54* and *Pdyn* (Fig 6L). Finally, pituitary expression of *Esr1* (Fig 6F; *p*=0.04, *d*=0.6), *Ar* (Fig 6G; *p*=0.05, *d*=0.6), *Cga*,(Fig 6H; *p*=<0.0001, *d*=1.4) and *Fshb* (Fig 6I; *p*=0.0001, *d*=1.2) levels were highest in animals with testes and independent of sex. A significant interaction between gonad and sex was seen when comparing means for *Lhb* (Fig 6J; *p*=0.01): males with testes had higher *Lhb* then females with testes (*p*=0.02, *d*=1.4) while males with ovaries had lower levels than males (*p*<0.0001, *d*=3.8) or females (*p*<0.01, *d*=2.1) with testes. Females with ovaries also had lower *Lhb* levels than males (*p*<0.0001, *d*=3.2) or females (*p*=0.03, *d*=1.6) with testes. *Lhb* did not significantly differ between females and males with ovaries (*p*=0.9, *d*=0.6). Pituitary *Gnrhr* expression was higher in males compared with females (Fig 6K; *p*=0.02, *d*=0.8) but independent of gonad type (*p*=0.07, *d*=0.6).

**Figure 6:**
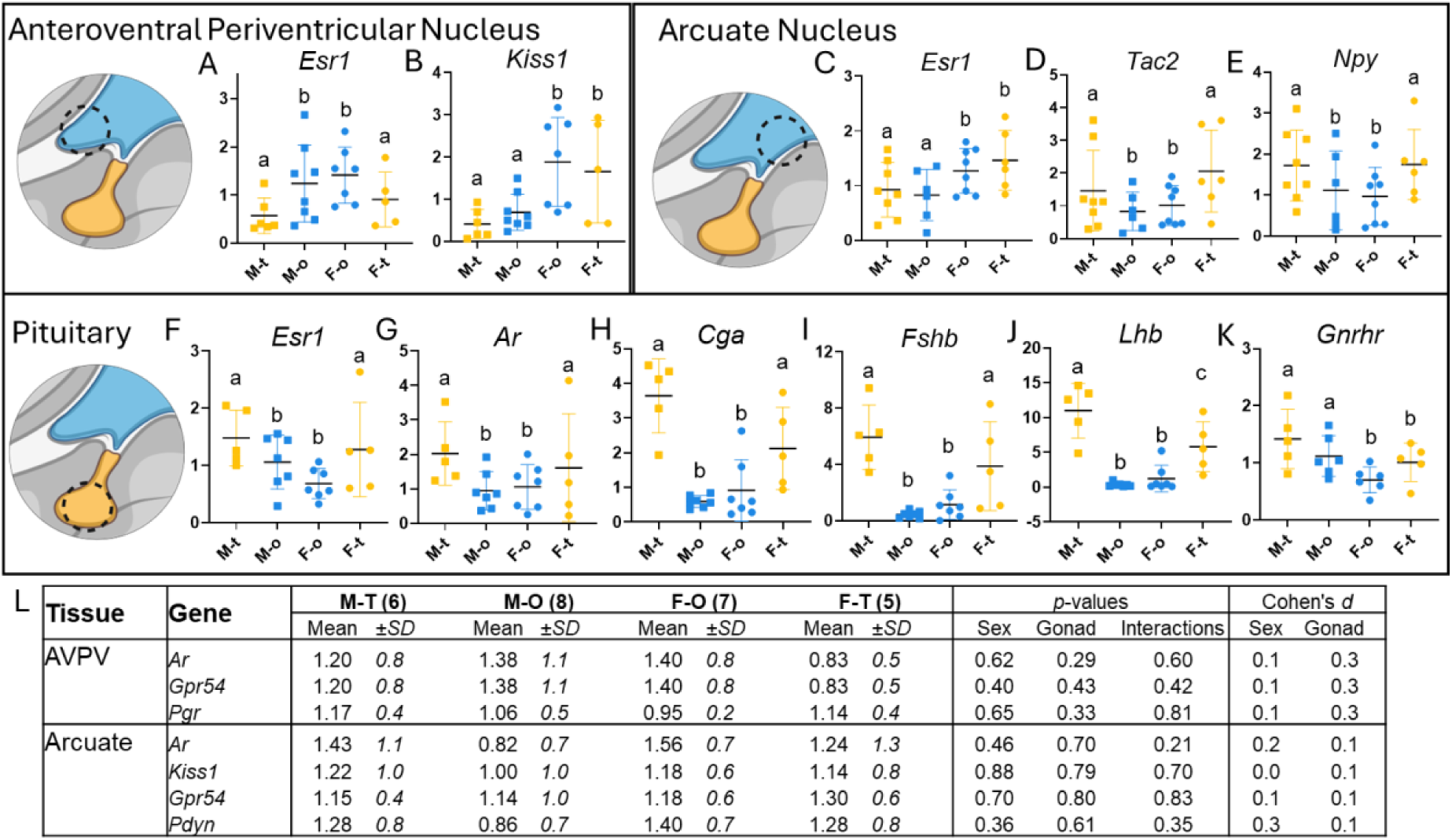
Brain and pituitary gene expression A) Animals with ovaries had higher *Esr1* expression in their anteroventral periventricular nucleus (AVPV) than those that received testes, regardless of sex. B) Conversely, *Kiss1* expression was highest in female animals compared with males, regardless of gonad type. C) Female animals had higher *Esr1* expression in their ARC than males, regardless of gonad type. Further, ARC). D) *Tac2* and E) *Npy* expression was highest in animals that received testes, regardless of sex. F-I) Animals with testes had the higher pituitary *Ar, Esr1, Cga,* and *Fshb* expression than those with ovaries, regardless of gonad type. J) Male animals with testes had the highest *Lhb* levels, followed by females with testes, while animals with ovaries had the lowest *Lhb* expression levels. K) Male animals had higher *Gnrhr* expression in their pituitary than females, regardless of gonad. L) No significant differences or large effects in AVPV *Ar, Gpr54*, and *Pgr* nor ARC *Ar, Kiss1, Gpr54*, and *Pdyn* expression were seen. Different letters are statistically different, *p*<0.05.

## Discussion

Here we show that the adult mouse hypothalamus and pituitary can integrate different-sex gonads into the host HPG axis. Multiple outcomes supported the incorporation of testis transplants. Though intact male mice initially had higher FSH than females, gonadectomy increased FSH levels, removed this sex difference. These high FSH levels likely supported the maturation of testis transplants^45,46^. Mature testis transplants could then respond to LH by producing T^47^ which provided negative feedback to the hypothalamus and pituitary to maintain HPG homeostasis^48^. Indeed, following transplantation of testes, FSH in males and females lowered to levels comparable to those of pre-gonadectomy males, suggesting reestablished negative feedback. Terminal steroid measures provided direct evidence of elevated T produced and secreted from the transplanted gonads. Further, terminal anatomy suggested sustained T elevation as androgen-sensitive tissues followed expected patterns for male^49,50^ and female mice^31^ with high T—males with testes had larger seminal vesicles than those with ovaries and females with testes displayed clitoromegaly. Histology revealed the presence of mature Sertoli and Leydig cells but limited spermatogenesis in testes transplants^51^. These elements are absent in testes from PD 6-9 pups^40,45,52^ and must have developed following transplantation. Leydig cells were located between seminiferous tubules, expressed receptors for LH, and were the likely source for elevated T in animals with testis transplants^47,53^. Though gametogenesis was not the focus of this study, the varying levels of sperm maturation seen in transplants required the presence of T^54–56^. Many seminiferous tubules had mature Sertoli cells, spermatogonia, and spermatocytes but no spermatids were seen. Given spermatogenesis is a 30-day cycle and requires Sertoli and Leydig cells to mature^57,58^, transplants may not have had sufficient time to complete spermatogenesis. Alternatively, processes may have inhibited spermatogenesis. Indeed, some tubules with spermatogonia had dilated lumens and lacked recognizable Sertoli cells. This morphology is similar to testes where spermatogenesis is prevented by altered physiology or disease states^59,60^. Notably, an increase in luminal pressure can lead to this morphology^61,62^ and encapsulation of transplanted gonads by host tissues may have increased pressure by blocking outflow from seminiferous tubules. Overall, we found that testis transplants addressed the needs of a novel method for providing gender-affirming hormone therapy; testis transplants in female mice matured, began producing T in response to host gonadotropins, and provided negative feedback to the hypothalamus and pituitary regardless of sex.

Functional integration of ovarian transplants was also supported by several findings. As before, gonadectomy elevated FSH levels in male and female animals prior to implantation and removed the sex difference. The ovaries respond to FSH by producing E, which communicates with the hypothalamus and pituitary in multiple ways throughout the gonad cycle^63–65^ to modulate gonadotropin release^19,66,67^. Ovary transplantation decreased FSH in males and females to levels comparable to females before gonadectomy. Further, female mice given ovaries had heavier uteri, which is highly responsive to E^68,69^. Interestingly, ovary transplants had no effect on terminal E levels in males or females. This may be due to the relatively low sensitivity of the E immunoassay used^70,71^. It will be critical to use more sensitive assays^72^ in future studies. Despite our inability to detect elevated E, we observed several E-dependent processes histologically. Ovaries taken from PD 6-9 pups contain mostly primordial and primary follicles^39^, which require both FSH and E to fully mature^73^. The appearance of primary, secondary, and antral follicles in ovary transplants indicates folliculogenesis occurred after transplantation. This process not only relies on E but involves E secretion from granulosa cells^64,74^. Overall, our findings suggest ovary transplants meet the needs of E-GAHT; ovaries transplanted into male mice matured through E-dependent processes and provided negative feedback to the hypothalamus and pituitary. In addition to E-producing follicles, transplanted ovaries from females and two from males had CLs. These LH-sensitive structures form from ovulated follicles and produce progesterone^75^, a critical component of E-GAHT for some patients^76^. An obvious barrier to using this mouse model is the rarity of progesterone-producing CLs in ovaries transplanted into male mice (n=2). Critically, our histological analysis may have failed to identify all CLs formed in ovary transplants as only half of the transplanted tissue was examined. In future studies, analysis of whole transplants will be important, allowing comparisons between potential subpopulations in males based on CL presence. The appearance of blood-filled cystic antral follicles in all ovary transplants taken from males, and many from females, are indicative of fully developed antral follicles that did not or could not ovulate^77^.

Cystic follicles may have matured shortly after transplantation in response to high FSH or even mechanical stimulation^64,73,78^, then reached maturity before fully integrating their positive feedback into the HPG. Once ovarian transplants could provide a signal sufficient to produce an LH surge^79^, antral follicles may have ovulated and formed CL. Indeed, males with CLs also had cystic follicles, so delaying terminal tissue collection may help bolster the number of CL-producing males in future research. To fully characterize the steroidogenic capacity of different-sex ovary transplants, it will be essential to examine relationships between CL formation, LH surges, and the local production of progesterone in future studies.

Although the adult human and primate HPG axis appears to possess sufficient plasticity to support different-sex gonad cycles^23,24,27^, it was assumed the HPG axis of rodents would be permanently organized by adulthood^80–84^. Our findings contrast these early assertions of either the functional limits or immutability of HPG sex differences in mice. Given gonad-dependent gonadotropin levels, we probed the expression of upstream HPG signaling genes. Genes that produce receptors or neurotransmitters related to gonadotropin release were investigated in two hypothalamic regions. The ARC contains a population of neurons that receive negative feedback from steroid hormones and express the neurotransmitters Kisspeptin (KISS), Neurokinin B (NKB), dynorphin, and neuropeptide y (NPY), called KNDy neurons. These neurotransmitters modulate the activity of both KNDy and gonadotropin-releasing hormone (GnRH) neurons to maintain HPG homeostasis^19,67^. Testis transplants increased *Tac2* (NKB) and *Npy* (NPY) mRNA, independent of sex. Increased NKB or NPY signaling may have elevated gonadotropin levels in animals with testes as genetic knockouts of NKB receptors are associated with gonadotropin deficiencies in humans and mice^85,86^ and NPY activates GnRH neurons^87^. Elevated NKB or NPY in animals with testes may have facilitated downstream changes in circulating gonadotropins^87,88^. Notably, the ARC is thought to produce LH pulses in males and females^89,90^. These changes may be necessary to ensure similar pulse generation under varying hormonal conditions. Indeed, the ARC functions similarly in males and females^89^ despite variability in Esr1 expression^91^. Interestingly, the sex difference in ARC *Esr1* expression was independent of gonad type. Notably, ARC Esr1 is also involved with non-HPG processes, such as calcium and energy homeostasis in bones^92^, so such changes may be unrelated to gonad-dependent HPG function.

Another steroid hormone sensitive HPG neuron population is the kisspeptin neurons in the AVPV. These also control GnRH neuron activity but receive both positive and negative feedback from the gonads leading to surges of LH. Two days of exogenous E treatments to mimic proestrus levels (E-priming) is sufficient positive feedback to induce an LH surge in female rodents, but not males^84^. Critically, AVPV neurons expressing *Esr1* are essential for producing LH surges^93^ and male mice typically have less AVPV *Esr1* mRNA than females^94^. These sex difference are thought to permit the production of LH surges in females^84^. Though E-priming fails to initiate an LH surge in males^94^, we saw evidence of LH surges via CL formation in ovaries from both males and females. Notably, E-priming elevates AVPV *Esr1* in females but not males^84^ yet we found ovary transplants increased *Esr1* expression in the AVPV, regardless of sex. This change in *Esr1* expression may rely on several mechanisms, such as changes in gene expression or even the integration of newborn neurons into the adult hypothalamus. Newborn cells are integrated into the AVPV throughout adulthood and blocking adult neurogenesis prevents the production of LH surges^38^. It is possible that established and/or newborn AVPV cells in male mice with ovaries took on characteristics normally found in females during prolonged exposure to ovaries. Two days of E-priming may be insufficient to shift AVPV function in males, but several weeks integrating signals from transplanted ovaries might do this. These findings call into question the rigidity of certain HPG sex differences, suggesting ongoing processes can change which signals activate organized circuits in adult mice. Conversely, we found expression of *Esr1* in the ARC and *Kiss1* in the AVPV is higher in females regardless of gonad. The retention of these sex differences suggests they may be dispensable for gonad-specific function and/or retain functional plasticity despite similar mRNA profiles.

Downstream from the hypothalamus, some mRNA expression levels in the pituitary aligned with high circulating gonadotropin levels in animals with testes. The subunits responsible for producing FSH and LH (*Fshb*, *Lhb*, and *Cga*) were highest in animals with testis transplants compared to those with ovaries. However, receptor gene expression appears to contradict gonadotropin outcomes. E can act directly on the pituitary to decrease gonadotropin release^95^ yet animals with ovaries had lower pituitary *Esr1* levels and gonadotropins than animals with testes. Similarly, GnRH binding to its receptor increases gonadotropins^66,96^ and males have higher *Gnrhr* expression overall compared with females but males with ovaries had lower gonadotropins than females with testes. This suggests feedback received by the hypothalamus is increasing gonadotropin subunit expression and circulating levels in animals with testes and/or lowering them in animals with ovaries. Though we have identified gonad-dependent gene expression in the ARC and AVPV, GnRH neurons are the direct link between these brain regions and the pituitary^66,87^. The diffuse distribution of these neurons could not be sampled using our methods but characterizing changes in GnRH neurons will be essential to fully understand gonad-dependent HPG function. Further, gonad differences in non-steroidal products, like inhibins^97^ and activins^98,99^, may act on and within these regions to influence gonadotropin release and/or gene expression. Overall, we identified gonad-dependent mRNA levels which may be essential for different-sex gonad incorporation into a novel HPG. Identifying the origins and functional significance of gonad-dependent outcomes may help understand and improve different-sex gonad transplants.

Beyond translation to gender-affirming care, gonad transplants in mice expand available *in vivo* research methods for examining sex differences^100^. The HPG feedback circuit produces cyclic steroidal and non-steroidal hormone levels, leading to differential activation of downstream pathways mediating innumerable measures of health^101,102^. All gonads can secrete E and T but hormone-dependent sex differences are currently investigated by manipulating main steroidal product of gonads or their receptors alone^103,104^. Gonad-dependent differences tied to health outcomes, like hypothalamic *Esr1* expression^105–108^, are obscured by the hormone replacement techniques used in sex difference research^109^. For example, AVPV *Esr1* levels are unaltered in gonadectomized males given E alone^84^ but we found elevated *Esr1* levels in gonadectomized males given ovaries. A same-and different-sex gonad transplant mouse model expands the sex characteristics available for manipulation in research, allowing basic and pre-clinical investigators to independently examine or compare gonad- and steroid hormone-dependent outcomes.

In conclusion, these data support the feasibility of different-sex gonad transplants in adult mice and their utility as a model for investigating this potentially novel method of providing gender-affirming hormones to TNG patients. Such a procedure would rely on donor tissue and require immune-suppressing or -isolating methods to function^110,111^. Mice may be used to understand and develop different-sex donor gonad transplants along with any necessary immunological interventions through controlled experimentation. An alternative strategy could be implantation of autologous stem-cell derived steroidogenic cells, produced through CRISPR/Cas9 manipulations and paracrine treatment of stem cells^112^. Characterizing health-related endpoints (musculoskeletal health glucose homeostasis, inflammatory pathways etc.) in future mouse experiments will also be necessary to inform the safety of gonad transplants, as will further studies in non-human primates with an eye toward eventual translation to humans. Finally, it is essential to consider how a novel gender-affirming technology will be received by the intended users^113,114^. In addition to investigating this model, researchers should seek collaborations with both transgender studies scholars^35,115^, to interrogate the implications of this technology, and TNG people^113,116^, to ensure study questions, interpretations, and products align with community needs and desires.

## Resource Availability

Data underlying this article will be shared on reasonable request to the corresponding author.

## Acknowledgements and Funding sources

We would like to thank members of the Shikanov lab for valuable feedback on manuscript figures and discussion. NIH T32 DK071212, R01 HD098233. Biorender.com was used to create figure icons.

## Author Contributions

Study Design (DRP, JGC, MAW, VP, AS, MBM), Acquisition (DRP, EC, JGC, RKC, REK, AS), Analysis (DRP, EC, JGC, REK, MAW, AS), Interpretation (All), Manuscript Drafting (DRP), Manuscript Preparation (All).

## Supplementary Tables

**Supplementary Table S1:**
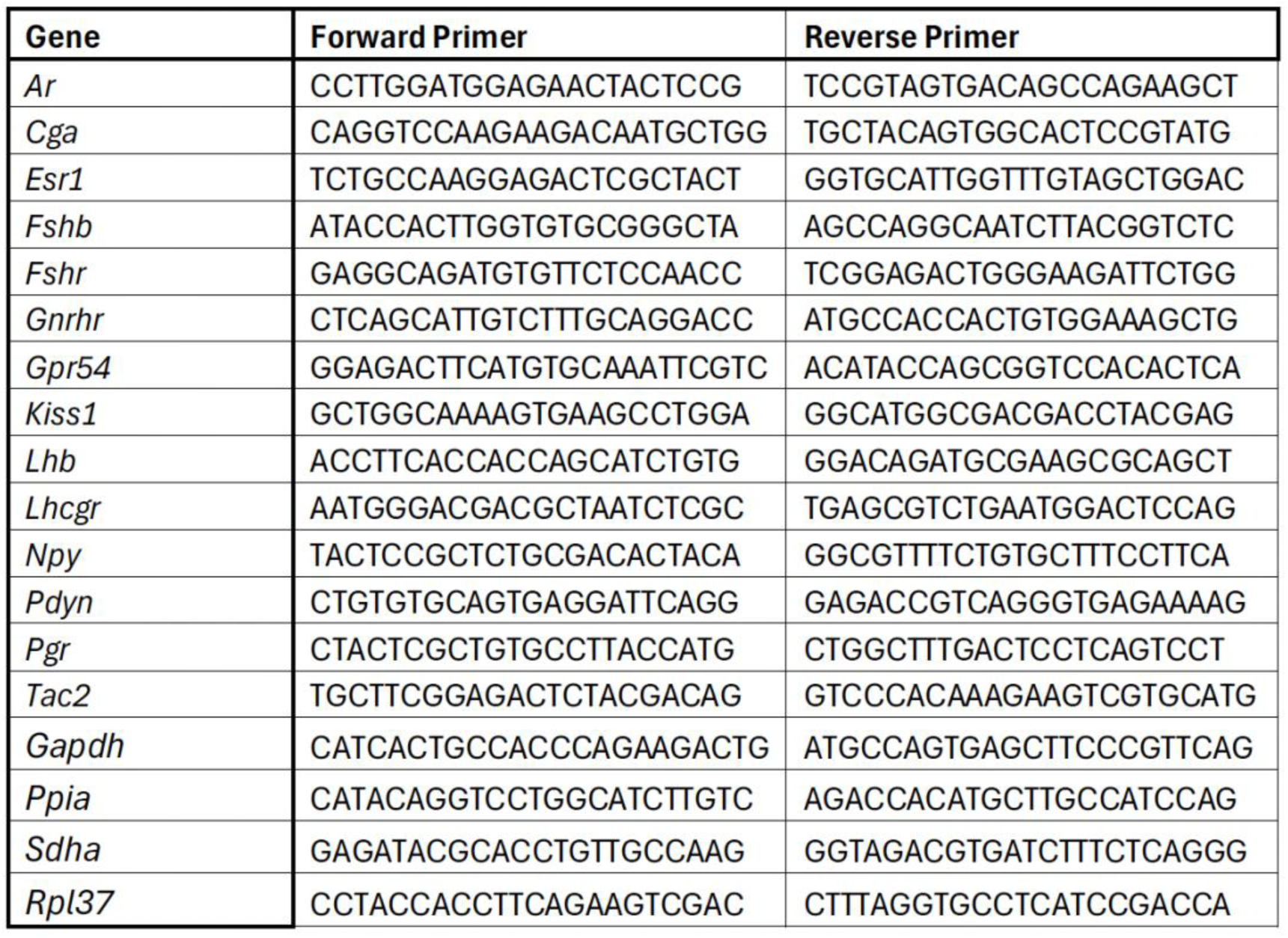
Primers for qPCR reactions.

**Supplementary Table S2:** Average hormone levels by group. M-t=Males with testis, M-o=Males with ovaries, F-o=Females with ovaries, F-t=Females with testis, FSH=Follicle stimulating hormone, LH=Luteinizing hormone.

| Hormone | Timepoint | M-t |  | M-o |  | F-o |  | F-t |  |
| --- | --- | --- | --- | --- | --- | --- | --- | --- | --- |
|  |  | Mean | ±SD | Mean | ±SD | Mean | ±SD | Mean | ±SD |
| FSH (ng/mL) | Initial | 38.27 | 6.5 | 37.91 | 9.4 | 5.71 | 5.3 | 2.65 | 0.1 |
|  | Post-GDX | 50.83 | 3.5 | 50.26 | 6.5 | 37.73 | 4.8 | 39.39 | 2.8 |
|  | 1 | 62.08 | 4.4 | 53.47 | 18.8 | 39.58 | 16.2 | 52.03 | 6.4 |
|  | 2 | 60.73 | 5.7 | 26.23 | 29.4 | 17.78 | 15.6 | 49.15 | 8.5 |
|  | 3 | 56.15 | 6.8 | 18.68 | 26.3 | 14.75 | 17.1 | 45.57 | 7.1 |
|  | 4 | 54.07 | 7.0 | 17.21 | 25.2 | 9.41 | 8.6 | 42.86 | 4.5 |
|  | 5 | 52.14 | 4.1 | 14.56 | 19.3 | 9.22 | 6.5 | 45.98 | 5.0 |
|  | Terminal | 41.70 | 7.2 | 15.37 | 21.9 | 16.36 | 12.7 | 41.79 | 8.2 |
| LH (ng/mL) | Terminal | 1.30 | 0.7 | 0.55 | 0.8 | 0.21 | 0.2 | 2.13 | 1.3 |
| Estradiol (pg/mL) | Terminal | 7.663 | 9.4 | 3.143 | 3.9 | 5.663 | 6.8 | 2.614 | 4.8 |
| Testosterone (ng/dL) | Terminal | 62.14 | 35.5 | 14.69 | 9.5 | 26.39 | 7.5 | 67.56 | 35.8 |

**Supplementary Table S3:**
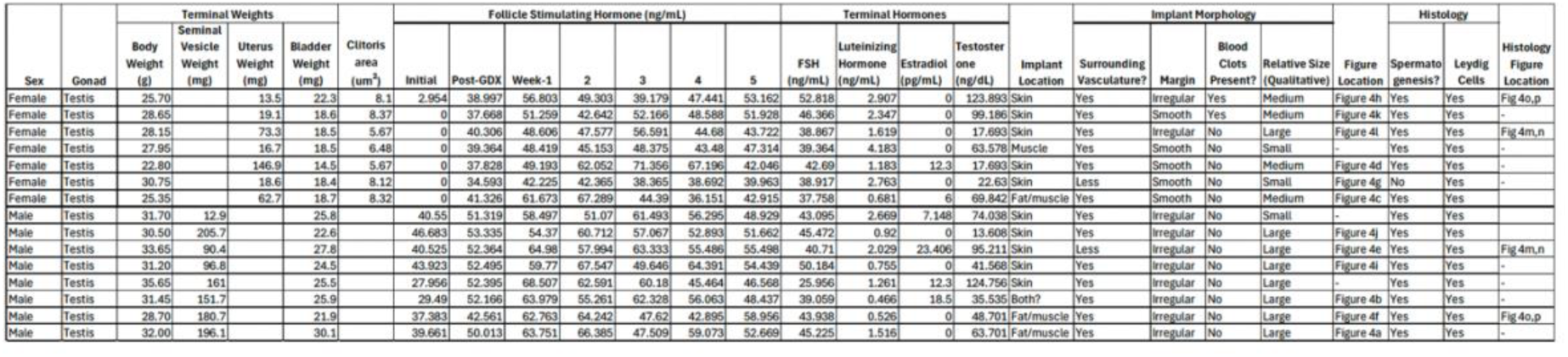
Testis transplant mouse data. GDX=gonadectomy, FSH=follicle stimulating hormone.

**Supplementary Table S4:**
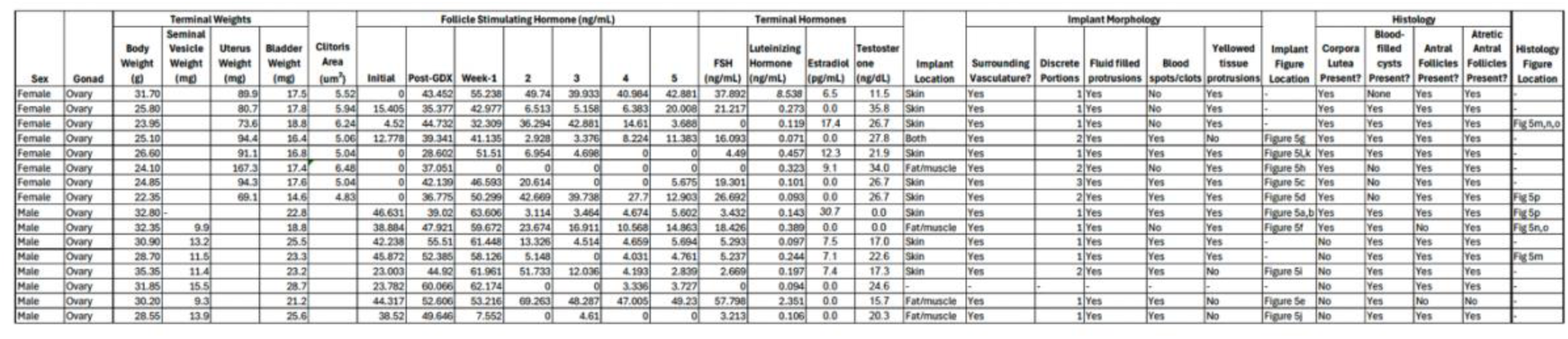
Ovary transplant mouse data. GDX=gonadectomy, FSH=follicle stimulating hormone.

